# Rho/ROCK activity tunes cell compartment segregation and differentiation in nephron-forming niches

**DOI:** 10.1101/2023.11.08.566308

**Authors:** John M. Viola, Jiageng Liu, Aria Huang, Samuel H. Grindel, Louis S. Prahl, Alex J. Hughes

## Abstract

Controlling the time and place of nephron formation *in vitro* would improve nephron density and connectivity in next-generation kidney replacement tissues. Recent developments in kidney organoid technology have paved the way to achieving self-sustaining nephrogenic niches *in vitro*. The physical and geometric structure of the niche are key control parameters in tissue engineering approaches. However, their relationship to nephron differentiation is unclear. Here we investigate the relationship between niche geometry, cell compartment mixing, and nephron differentiation by targeting the Rho/ROCK pathway, a master regulator of the actin cytoskeleton. We find that the ROCK inhibitor Y-27632 increases mixing between nephron progenitor and stromal compartments in native mouse embryonic kidney niches, and also increases nephrogenesis. Similar increases are also seen in reductionist mouse primary cell and human induced pluripotent stem cell (iPSC)-derived organoids perturbed by Y-27632, dependent on the presence of stromal cells. Our data indicate that niche organization is a determinant of nephron formation rate, bringing renewed focus to the spatial context of cell-cell interactions in kidney tissue engineering efforts.

## Introduction

Nephrons created during embryonic development of the kidney confer its fundamental filtration role during adult life. Variability in nephron number (‘endowment’) is high even among normal kidneys, and is exacerbated by certain congenital defects. This has a clinical impact because low nephron endowment correlates with increased risk of hypertension and chronic kidney disease (Hoy et al. 2005; Bueters, van de Kar, and Schreuder 2013; Perl, Schuh, and Kopan 2022; Yermalovich et al. 2019; Keller et al. 2003). Kidney transplant remains the only opportunity for long-term functional replacement in humans, but donor organs are in short supply (Matas et al. 2015). A better understanding of cell interactions and decision-making in nephron progenitors that compound to set nephron endowment would benefit efforts to both therapeutically manipulate nephron number and to construct them *de novo*.

The developing kidney cortex is filled with nephron progenitor stem cell niches (the ‘metanephric/cap mesenchyme’ and its interfaces with surrounding cell compartments) that create nephrons throughout kidney development. Each cap interfaces with the surface of a branching ureteric bud tip (the future urinary collecting ducts), and with stromal cells at remaining interfaces (**Fig. 1A,B**). These three cell compartments exchange paracrine and juxtacrine cues, suggesting that the geometry and contact area among them may play a role in nephron progenitor cell decision-making. Recent advancements in cell patterning and bioprinting technologies are promising approaches to building tissue niches *in vitro* (Todhunter et al. 2015; Viola et al. 2020; Skylar-Scott et al. 2022). Closing knowledge gaps in the relationship between nephron progenitor niche structure and nephron formation would allow researchers to harness the power of cell patterning techniques to drive functional kidney tissue engineering outcomes *in vitro* (Combes et al. 2016; Lefevre et al. 2017; Oxburgh et al. 2004; Homan et al. 2019; Lawlor et al. 2021, 2019).

**Fig. 1:**
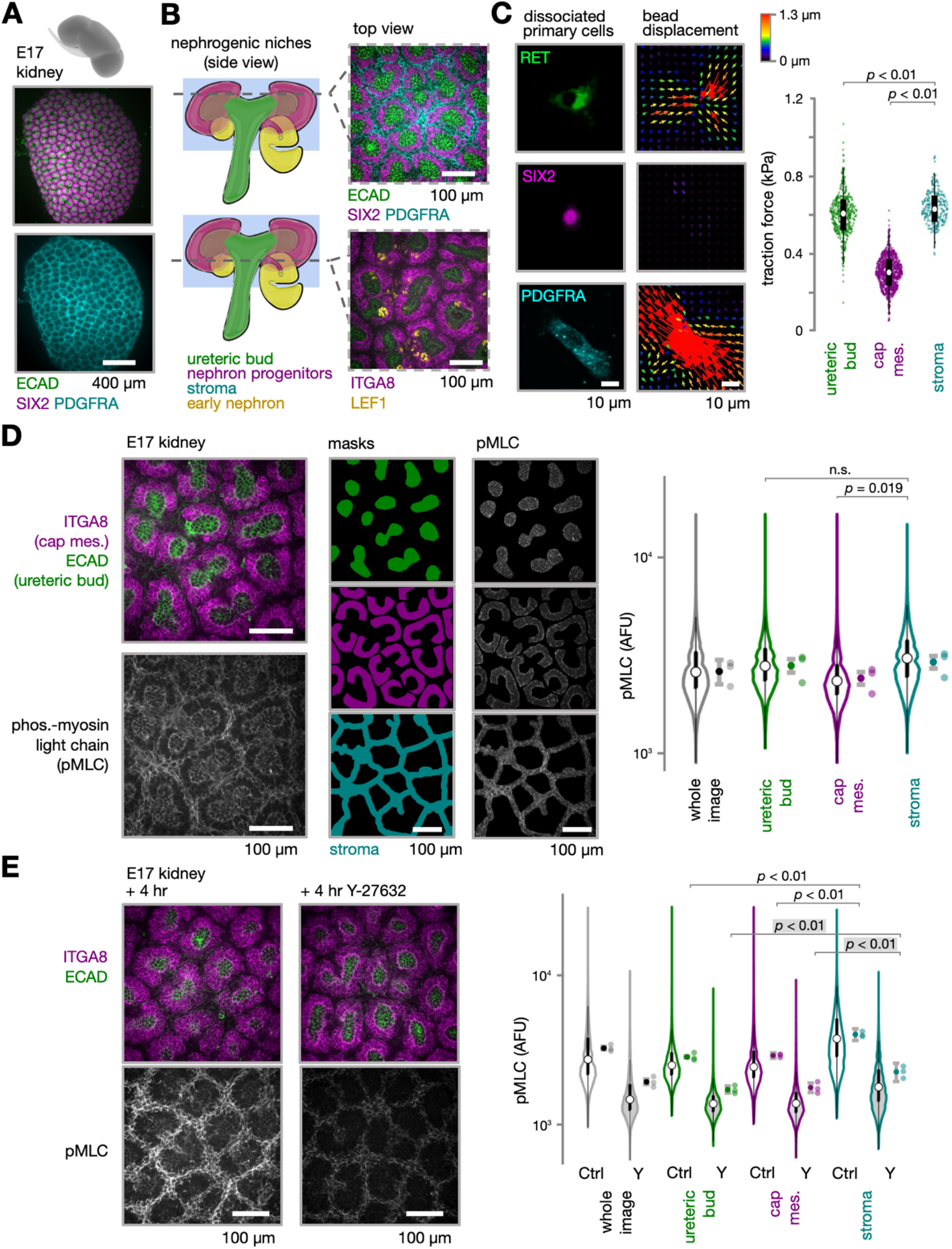
Traction force microscopy and pMLC staining indicate a mechanical tension hierarchy in the mouse nephrogenic niche that is homogenized by ROCK inhibition. **(A)** Embryonic day 17 (E17) mouse kidney cortex immunofluorescence micrographs showing arrayed nephrogenic niches. (**B**) *Left,* Schematic of niche anatomy side view. Dotted lines approximate optical depth of immunofluorescence microscopy images. *Right,* confocal fluorescence micrographs at indicated imaging depths. (**C**) *Left*, Confocal fluorescence micrographs of example primary mouse embryonic kidney ureteric bud epithelial, nephron progenitor, and stromal cells on polyacrylamide TFM substrate and corresponding traction force heatmaps. *Right*, violin plot of traction force by niche cell type (mean ± S.D., *n* > 282 cells per condition, ANOVA with Tukey’s test *p* values indicated) (**D**) Left, Whole-mount confocal fluorescence micrographs of E17 kidney niches, pMLC channel isolated at bottom. *Middle*, Manually segmented niche cell compartment masks and corresponding pMLC channel masked by cell compartment. *Right*, Violin plots of pMLC pixel arbitrary fluorescence (AFU) in the example image by cell compartment (median and quartiles), and means of similar fields of view fields of view (*n* = 3 fields of view, paired *t*-test *p* values indicated). (**E**) Similar images and plots for kidney explants cultured for 4 hrs ± 30 µM Y-27632. Images were collected at equivalent settings.

Cell sorting into tissue compartments depends on differences in contractility and adhesion (Steinberg 1963; Cerchiari et al. 2015; Tsai et al. 2020). In fact, spontaneous cell sorting outcomes of primary kidney cell mixtures can be disrupted or enhanced by perturbing the activity of the actin cytoskeleton (Leclerc and Costantini 2016; Meyer et al. 2006). Additionally, regulation of Rho GTPase, Rho-associated protein kinase (ROCK), and non-muscle myosin IIA/B isoforms (Myh9/Myh10) in the developing kidney can increase nephron formation (e.g. low dose H1152, a ROCK inhibitor) or decrease it (e.g. high dose H1152; mesenchymal Myh9/Myh10 deletions)(Lindström, Hohenstein, and Davies 2013; Recuenco et al. 2015; Tanigawa et al. 2015). One hypothesis for these differential outcomes is that tissue structural context (cell sorting and migration) determined by cell biophysical properties (adhesions, tensions) is an underlying regulator of niche signaling, including nephron progenitor cell exposure or sensitivity to biochemical cues.

There are several examples of such interactions between niche structure and signaling. Canonical Wnt signaling in nephron progenitors, driven by ureteric bud-derived ligands, plays an important role in nephron progenitor decision-making. Low levels of Wnt signaling promote maintenance of the progenitor pool while high levels promote differentiation (Park, Valerius, and McMahon 2007; Park et al. 2012). However, other signaling pathways including those modulated by ligands derived from the stromal compartment have been shown to influence nephron progenitor decision-making as well (Fetting et al. 2014; Ohmori et al. 2015; Rowan et al. 2018; Das et al. 2013). Stromal Fat4 expression, a cell-cell contact-dependent ligand, enhances phosphorylation and cytoplasmic retention of the transcription factor YAP in nephron progenitors, tipping their perception of Wnt/β-catenin signaling toward differentiation (Das et al. 2013). In fact, stromal knockout mouse embryonic kidneys have a severe defect in nephron formation and contain oversized nephron progenitor pools (Hum et al. 2014). Additionally, decreasing nephron progenitor attachment to the ureteric bud by genetic knockout of Wnt11 increases progenitor cell proximity to stromal cells and correlates with increased nephron formation events (O’Brien et al. 2018).

Here, we find that decreasing cell tension primarily in the stroma of the developing kidney cortex through brief ROCK inhibition increases mixing between niche cell compartments and non-autonomously promotes differentiation of nephron progenitors. The increased nephron differentiation effects of Y-27632 are seen only in the presence of stromal cells, suggesting a relationship between niche cell biophysical properties, niche organization, stromal-nephron progenitor contact, and renewal vs. differentiation decision-making. We also show that Y-27632 increases nephron differentiation events in human iPSC-derived kidney organoids. Our data suggest that successful *ex vivo* nephrogenesis through bioprinting and other cell patterning technologies should carefully consider niche cell type ratios and geometry.

## Results

### Cell tension differences between nephrogenic niche compartments drives niche structural integrity

We hypothesized that the ratio of interfacial adhesion:tension among cell compartments may contribute to the organization of nephron progenitor niches (Combes, Davies, and Little 2015). We first assessed differences in the contractile properties of the ureteric bud, cap mesenchyme, and stromal cell types constituting the niche using traction force microscopy (TFM) (Tseng et al. 2012). We recently combined TFM with immunofluorescence to retrieve cell identities and traction forces (Hughes Lab, *In preparation*). Dissociated cells from embryonic day (E)17 mouse kidneys showed clear differences in traction force among niche cell compartments (**Fig. 1C**). The stromal cells that line each nephrogenic niche exerted the highest traction force, followed closely by the ureteric bud cells, while the nephron progenitor cells exerted roughly half the traction force as the other two populations. These data predict that stromal and ureteric bud cells are dominant contributors to interfacial tension in the nephrogenic niche. We next quantified phosphorylated myosin light chain (pMLC) in these niche compartments by confocal immunofluorescence, a correlate of myosin activation and cytoskeleton contractility (Parada et al. 2022; Shiraishi et al. 2023) (**Fig. 1D**). We analyzed an optical slice that passes through the top (cortical)-most portion of the ureteric bud to capture the ureteric bud, cap mesenchyme and stroma in the same image. We found that the stromal compartments in E17 kidneys had significantly higher pMLC signal, followed by the ureteric bud and nephron progenitor compartments, corroborating the TFM data. Together these data reveal a mechanical tension hierarchy in the nephrogenic niche.

We next wondered if this tension hierarchy is necessary for niche organization, as a prelude to correlating the latter with nephron formation. To explore this, we targeted the Rho/ROCK pathway, a master regulator of the actin cytoskeleton (Maekawa et al. 1999; M. Amano et al. 1997). Rho/ROCK activity is required for normal nephron organization and tubule segmentation later in nephrogenesis (Lindström, Hohenstein, and Davies 2013). However, its effect on native niche organization and early nephron progenitor cell decision-making has not been closely studied. We performed brief (4 hr) explant culture of E17 mouse embryonic kidneys with the ROCK inhibitor Y-27632, which competes with ATP for binding to the catalytic site of ROCK, reducing actomyosin contractility, stress fiber and focal adhesion assembly (Ishizaki et al. 2000). Y-27632 caused a decrease in pMLC immunofluorescence signal in all compartments, and most dramatically in the stroma (**Fig. 1E**). The overall effect here was greater homogeneity in pMLC signal between compartments (stroma:ureteric bud pMLC ratio dropped from 1.58 to 1.45, and stroma:cap mesenchyme pMLC ratio dropped from 1.45 to 1.32). This accompanied a loss of spatial definition of stromal boundaries between niches. Specifically, SIX2+ cap mesenchyme progenitors were more frequently observed between niches; mixed in among the stromal cells at niche boundaries. Such nephron progenitor excursions into the stroma and even to neighboring niches has occasionally been observed in untreated embryonic kidney explant cultures (Combes et al. 2016). These ‘wayward’ cells contributed to spikes in average SIX2 immunofluorescence signal intensity across the niche gap (**Fig. 2A**). We then aligned and plotted the average and standard deviation of the SIX2 signal intensity profiles across a sample of gaps (**Fig. 2B**). We found that the average standard deviation in fluorescence intensity along the SIX2 fluorescence profile was greater in Y-27632-treated kidneys (**Fig. 2B**) likely due to decreased “smoothness” in the Y-27632 SIX2 profiles as seen in **Fig. 2A**. We fitted a sum of two Gaussian distributions to the average SIX2 fluorescence, revealing a modest and trending increase in niche width in Y-27632 treated samples. Both observations indicate increased spreading of the cap mesenchyme compartment after Y-27632 treatment. Similarly for the stromal compartment, we observed a modest 1.7 µm increase in width of Gaussian fits to the plot of the average PDGFRA signal across niche gaps (**Fig. 2C**). Since the cap mesenchyme and stroma compartments were both broader we reasoned that there was likely more mixing between the two compartments (**Fig. 2D**). We reasoned that this would be reflected in increased overlap between the compartments in our immunofluorescence images. We thresholded SIX2 and PDGFRA fluorescence to generate binary images of each compartment and used “AND” logic to keep only pixels located in both compartments (**Fig. 2E**). Indeed, we observed a ∼25% increase in pixels positive for both compartments after Y-27632 treatment, indicating a net increase in nephron progenitor-stromal mixing. We saw a similar effect qualitatively using the nuclear stromal marker Meis1/2 across optical planes spanning the cap mesenchyme (**Fig. S1**). In contrast, ureteric bud-cap mesenchyme interfaces remained intact, perhaps due to a ‘fail-safe’ mechanism of cell-cell adhesion associated with ureteric bud apicobasal polarity (Leung and Brugge 2012; Packard et al. 2013; Combes, Davies, and Little 2015). Together these data validate tissue-scale effects of Y-27632, namely a loss of niche boundary integrity, increased nephron progenitor dispersion, and increased probability of heterotypic contacts between nephron progenitors and stromal cells when traction and pMLC markers of cell collective tension are reduced using Y-27632.

**Fig. 2:**
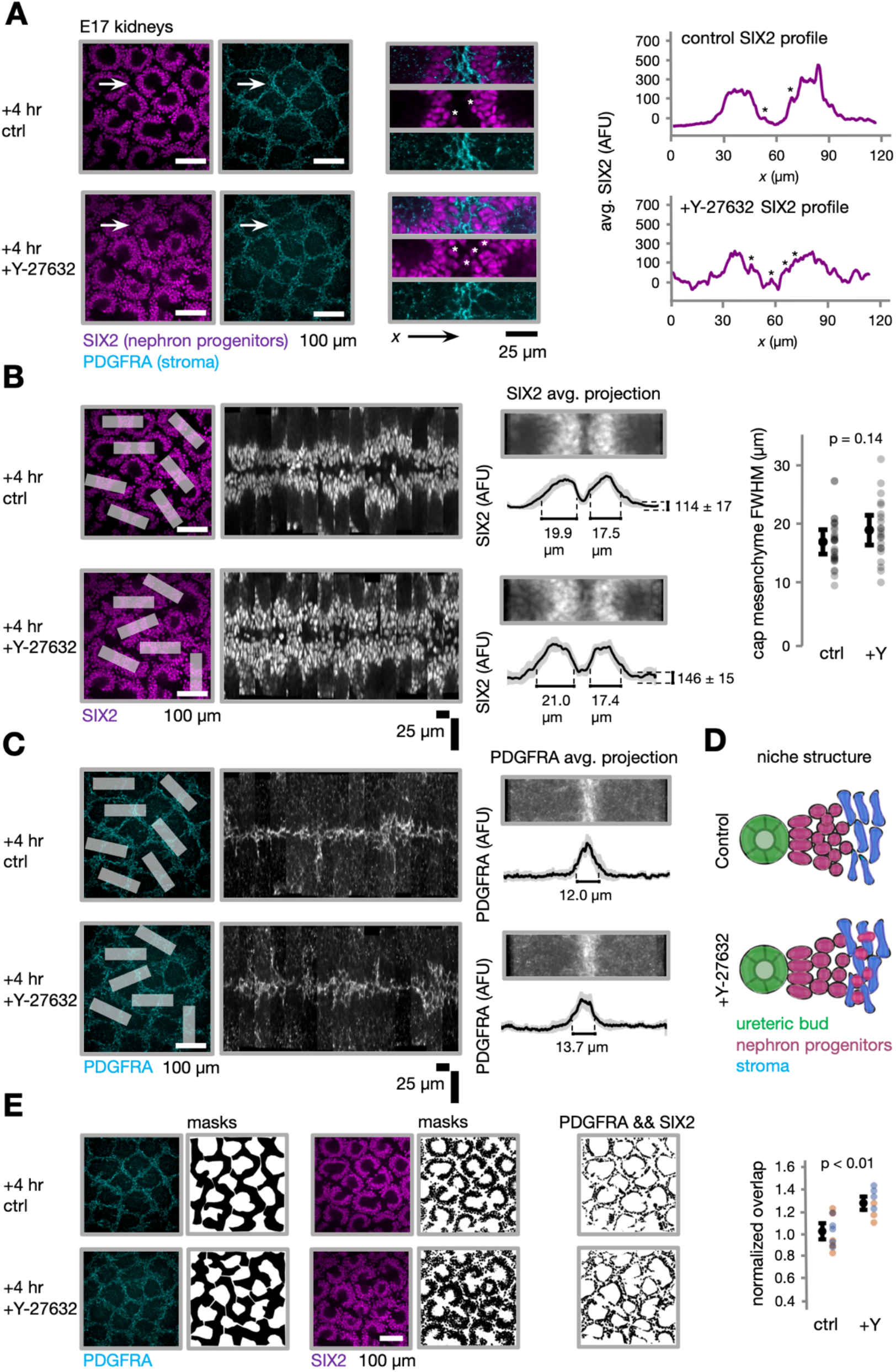
ROCK inhibition causes niche disorganization by mixing between nephron progenitor and stromal cell compartments. (**A**) *Left*, Confocal immunofluorescence micrographs of E17 niches after 4 hr culture in control media (top row) or 30 µM Y-27632 (bottom row). *Insets*: detail of regions indicated by white arrows. *Right*, Spatial fluorescence traces on SIX2 (nephron progenitor) channel. Black stars indicate contribution of individual nephron progenitors at or within the stromal boundary. (**B**) *Left*, immunofluorescence micrographs of E17 niches with some example stromal “gap” regions highlighted used in further analysis. *Middle,* Montages of individual “gap” regions of interest (ROIs) spatially registered to align their stromal boundaries along a common centerline. *Right*, Average projection images across ROIs and spatial fluorescence traces on SIX2 channel. Plot shows full width at half maximum (2.355*sigma) of Gaussian fits for ± Y-27632 niches (mean ± S.D., n > 22 niches per condition, *t*-test *p* value indicated). (**C**) Similar images and plots for PDGFRA (stroma) channel. **(D)** Schematic of nephron progenitor and stroma compartment mixing phenotype. (**E**) *Left*, PDGFRA and SIX2 immunofluorescence images of example E17 niches, associated segmentation masks, and image resulting from ‘AND’ operation (black pixels indicate channel co-localization). *Right*, nephron progenitor:stromal compartment mixing plotted as relative proportion of co-localized pixels for Y-27632-treated explants vs. controls (mean ± S.D., n > 7 images across 3 kidneys from 2 separate litters per condition, *t*-test *p* value indicated). Orange and blue data points denote images taken from kidneys of separate litters.

### Y-27632 promotes nephrogenesis in mouse kidney explants

Since stroma is generally thought to be pro-nephrogenic, we next hypothesized that the increase in nephron progenitor-stromal mixing induced by Y-27632 could impact nephron progenitor differentiation. We used long-term culture of E13 embryonic kidney explants to assess this in native niches. Since culturing the embryonic kidney in a fully 3D *ex vivo* context has been technically challenging in the field (Rosines et al. 2010; Sebinger et al. 2010), we turned to an established air-liquid interface culture system in which embryonic kidneys continue to undergo branching morphogenesis and nephrogenesis for at least 3-5 days (Lindström et al. 2015). Previous work has shown that long-term ROCK inhibitor treatment has concentration-dependent effects on branching of the ureteric bud *ex vivo*, and overall size and growth of cultured kidneys (Meyer et al. 2006; Lindström, Hohenstein, and Davies 2013). To avoid effects on branching and overall organ growth while still perturbing niche organization, we employed a brief (4 hr) pulse of ROCK inhibition (30 µM Y-27632) in E13 embryonic kidney cultures followed by a further 24 hr of culture in the control medium. While volume was unaffected (**Fig. S2**), we observed that Y-27632 increased explant surface area under the action of the air-liquid surface tension within 4 hrs, indicating a reduction in cell collective tension/stiffness (**Fig. 3A**). Y-27632 treatment also caused similar changes in niche organization as for the E17 kidneys shown in Fig. 2, namely increased dispersion of the cap mesenchyme compartment (**Fig. 3B**). We annotated JAG1+ nephron structures 24 hr after the removal of Y-27632 (**Fig. 3C,D**), allowing enough time for treated nephron progenitors to form nephron structures. We measured a modest increase in nephron number per kidney explant in addition to a significant 14% increase in nephron number per unit volume in Y-27632-treated explants vs. untreated controls (**Fig. 3E**). These data indicate that brief Y-27632 treatment was sufficient to change explant mechanics, niche organization and increase nephrogenesis in embryonic kidney explants.

**Fig. 3:**
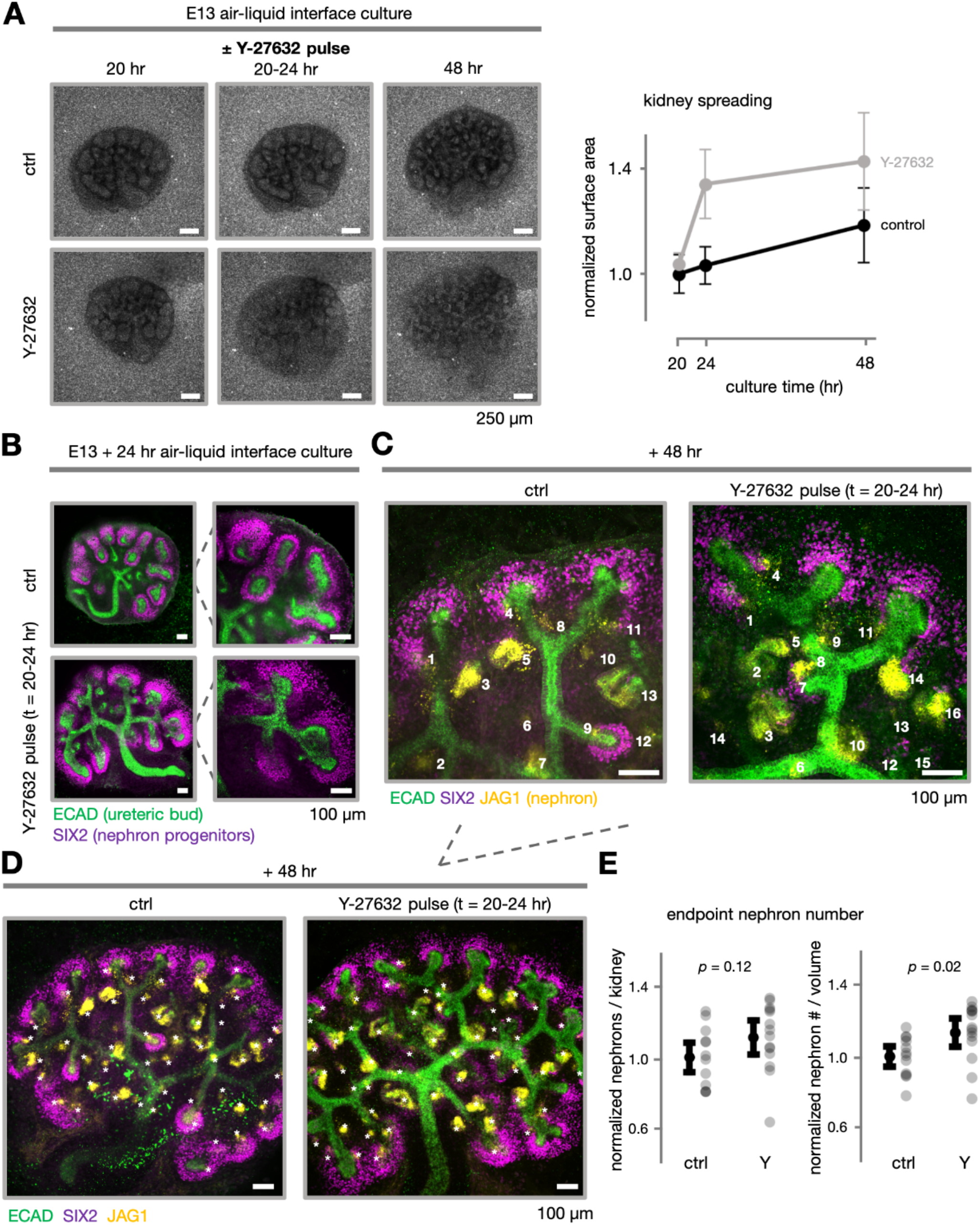
Brief ROCK inhibition increases nephrogenesis in mouse embryonic kidney explant cultures. (**A**) *Left*, Brightfield micrographs of E13 mouse embryonic explant cultures imaged through transwell membranes over culture period ± 30 µM Y-27632 pulse at t = 20-24 hr post culture. *Right*, plot of explant area growth over time, normalized by initial average control explant area (mean ± S.D., n > 8 kidneys across 2 litters per condition). (**B**) Confocal whole-mount immunofluorescence micrographs and insets showing detail of altered niche organization after 4 hr Y-27632 pulse. **(C)** Sample annotation of JAG1+ nephrons at 20x magnification. **(D)** Sample annotation of Jag1+ structures in representative whole explants at experiment endpoint. (**E**) Plots of nephron number and nephron density after explant culture ± Y-27632 normalized within litter (mean ± S.D., n > 12 kidneys per condition across 4 litters, unpaired *t*-test *p* values indicated).

### Y-27632 promotes nephrogenesis only in the presence of stroma in mouse kidney primary cell spheroids

We next sought to determine if the observed increase in nephron differentiation driven by Y-27632 was caused by a change in nephron progenitor-stromal cell interaction. We turned to a reductionist approach to generate mixed primary nephron progenitor/stromal cell spheroids from mouse embryonic kidneys (Brown et al. 2011). Nephron formation is induced in these spheroids by mimicking ureteric bud-derived Wnt signaling using the GSK3β inhibitor CHIR99021, but in the absence of additional variables associated with the presence of ureteric bud and branching morphogenesis. Embryonic mouse kidneys are briefly dissociated to recover cortical ‘nephrogenic zone’ (cortical layer) cells, which are reported to contain about 60% nephron progenitors and 15% stromal populations. The remaining cells are a mix of endothelial progenitors, mature nephron, ureteric bud, or immune cells (Brown et al. 2013). We validated enrichment for E17 kidney niche cells using marker immunofluorescence (**Fig. 4A**). Further magnetic enrichment has been reported to yield spheroids with >90% nephron progenitors, enabling a comparison between nephrogenic spheroid cultures containing or lacking stromal cells. We used similar induction conditions as described for human iPSC-derived nephron progenitor organoid differentiation (M. Takasato et al. 2014; Minoru Takasato et al. 2015; Morizane et al. 2015; Taguchi et al. 2014), namely a 3 µM CHIR pulse and ongoing 10 ng ml^-1^ fibroblast growth factor 9 (FGF9) treatment, and compared differentiation responses of primary cell spheroids with or without perturbation of cells with 10 µM Y-27632 during their initial aggregation. We then removed the CHIR and Y-27632 treatments and cultured the spheroids in basal media for another 24 hr. Y-27632 caused a significant increase in LHX1+ early nephron structures (a pre-tubular aggregate/renal vesicle marker) in mixed NPC-stromal spheroids, but not in purified NPC spheroids (**Fig. 4B**). Y-27632 has been shown to increase viability of stem cells in suspension and aggregation efficiency/kinetics (Watanabe et al. 2007; Ungrin et al. 2008). To eliminate these potential confounding effects and to test the effect of more conservative Y-27632 treatment, we repeated the same treatments on spheroids pre-aggregated in basal media, and for a shorter 4 hr pulse treatment of 3 µM CHIR +/- 10 µM Y-27632 (**Fig. 4C**). We again saw a significant increase in LHX1+ nephron structures in NPC-stroma spheroids after Y-27632 exposure, but the opposite effect in enriched nephron progenitor spheroids. This suggests that rather than being intrinsic to nephron progenitors, the increase in nephron differentiation in response to Y-27632 requires the presence of stromal cells and is independent of cell viability or aggregation effects.

**Fig. 4:**
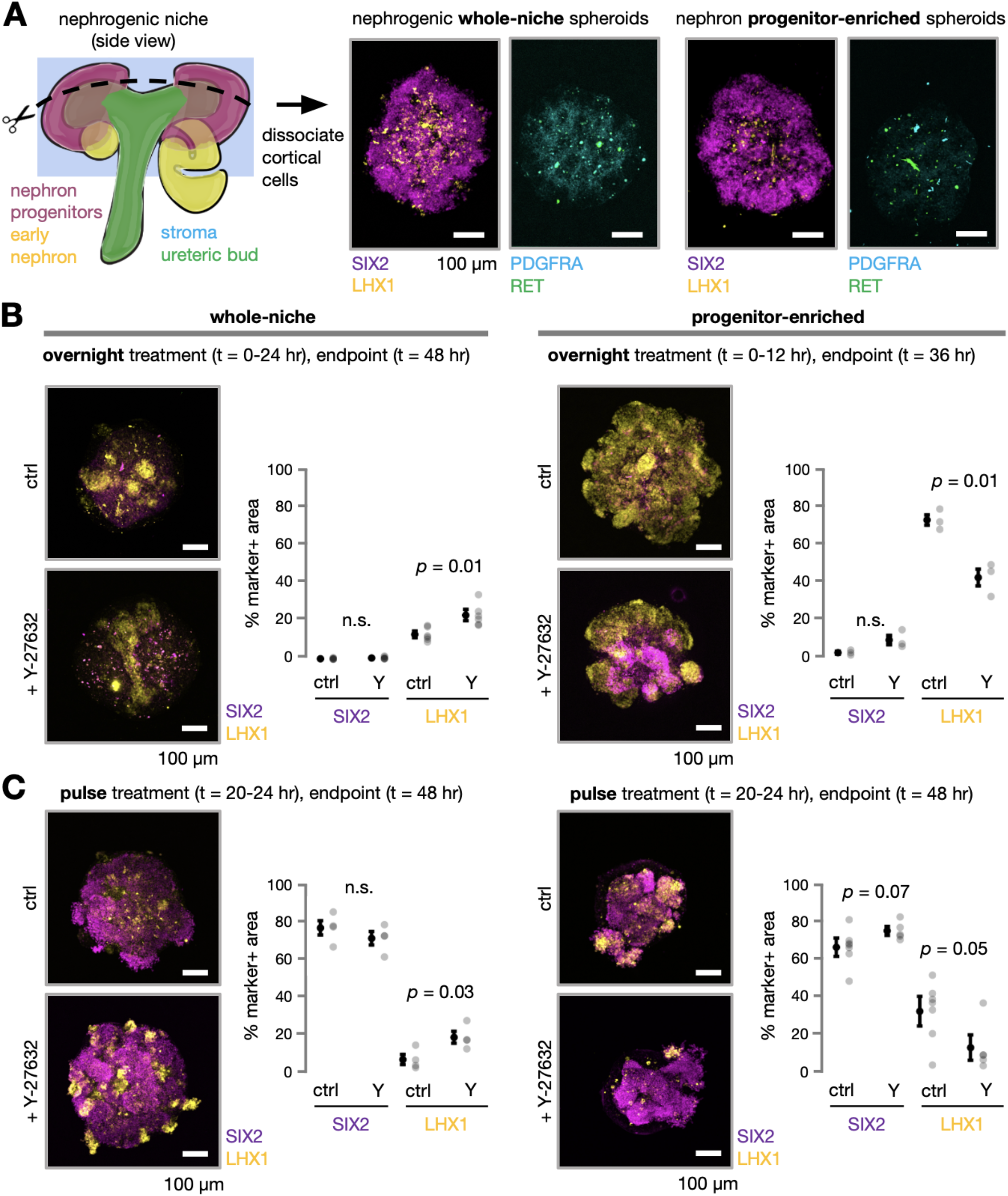
Brief ROCK inhibition increases nephrogenesis in mouse embryonic kidney primary cell spheroids, but only in presence of stroma. (**A**) *Left,* Schematic showing approximate dissociated region of E17 kidney cortex for primary ‘nephrogenic zone’ cell collection. *Right,* Confocal immunofluorescence micrographs of representative spheroids derived after primary cell aggregation, with or without further enrichment for nephron progenitors by magnetic negative cell sorting. (**B**) Representative immunofluorescence micrographs and spheroid composition plots at differentiation endpoint after overnight 3 µM CHIR ± 10 µM Y-27632 treatment for whole-niche nephrogenic zone cell spheroids, *left*, and nephron progenitor cell-enriched spheroids, *right* (mean ± S.D., *n* > 3 spheroids per condition, unpaired *t*-test *p* values indicated). (**C**) Similar images and plots for a 4 hr 3 µM CHIR treatment ± 10 µM Y-27632 pulse rather than overnight treatment.

### Y-27632 promotes nephrogenesis in human iPSC-derived kidney organoids

We next wondered if our observations in mouse primary cell spheroids would translate to human iPSC-derived kidney organoids, relevant to increasing their nephrogenesis efficiency. When differentiated, these roughly mimic the progression in transcriptional states and marker expression profiles of mouse nephron progenitors *in vivo*, despite some species-specific differences (Morizane et al. 2015; Minoru Takasato et al. 2015; Combes et al. 2019; Lindström et al. 2018; Wu et al. 2018). Human iPSC-derived kidney organoids contain a mixture of nephron progenitors and stromal cell populations, the latter congruous with human fetal kidney stroma despite missing some sub-populations and containing other off-target cells (Wilson and Little 2021). Before testing perturbations to Rho/ROCK signaling during nephron progenitor commitment, we first differentiated iPSCs through late primitive streak and posterior intermediate mesoderm (PIM) lineages to metanephric mesenchyme (nephron progenitors) according to established protocols (Morizane and Bonventre 2017), confirming expression of differentiation markers by qPCR (**Fig. S3**). We also validated that induced metanephric mesenchyme cells go on to form later nephron structures including ECAD+ tubules and cells expressing the proximal tubule marker LTL, the glomerular podocyte marker NPHS1, or the loop of Henle marker SLC12A1 (Na^+^/K^-^/2Cl^-^ cotransporter) after 21 days in culture (**Fig. 5A,B**). These structures are reported to be surrounded by MEIS1+ stroma (Porter et al. 2023). We next dissociated day 9 induced metanephric mesenchyme cells and plated them into low-attachment round-bottom plates, where they formed organoids that we assayed for early nephrogenesis. Similar to mouse primary cell spheroid experiments, we combined a 48 hr 3 µM CHIR pulse as in Takasato *et al*. (Minoru Takasato et al. 2015) with 10 µM Y-27632 or 30µm blebbistatin, a non-muscle myosin II ATPase inhibitor. We quantified tension differences in control, Y-27632 treated, and blebbistatin treated organoids using a laser ablation approach and found that Y-27632 significantly decreases organoid rebound velocity after ablation by ∼80%, while blebbistatin caused a small trending decrease (**Fig. 5C**). Interestingly, Y-27632 treated spheroids had similar rebound velocities (inferred tension) as nephron-forming niches in the E17 mouse kidney cortex (Viola et al. 2023). Organoids were analyzed immediately after the 48 hr treatment to catch early differentiation events by immunofluorescence for SIX2 and the early nephron marker LHX1 (**Fig. 5D**). Combining CHIR with Y-27632 induced a synergistic pro-differentiation effect. LHX1+ cells clustered in Y-27632-treated organoids and began to express the later medial nephron marker Jag1, indicating appropriate progression of nephron differentiation. Similar to our mouse primary spheroid assays, these data together show that Wnt-driven nephron progenitor commitment in human kidney organoids is higher when combined with Y-27632. The fact that a similar effect is observed for blebbistatin suggests that the effect of Y-27632 is likely being mediated at the level of myosin activity/cell collective tension rather than other signaling arms downstream of Rho/ROCK (Mutsuki Amano, Nakayama, and Kaibuchi 2010). These findings indicate a potential parallel between nephrogenesis in human organoids and our findings relating niche cell compartment contractility, cell sorting, and nephron differentiation in the mouse.

**Fig. 5:**
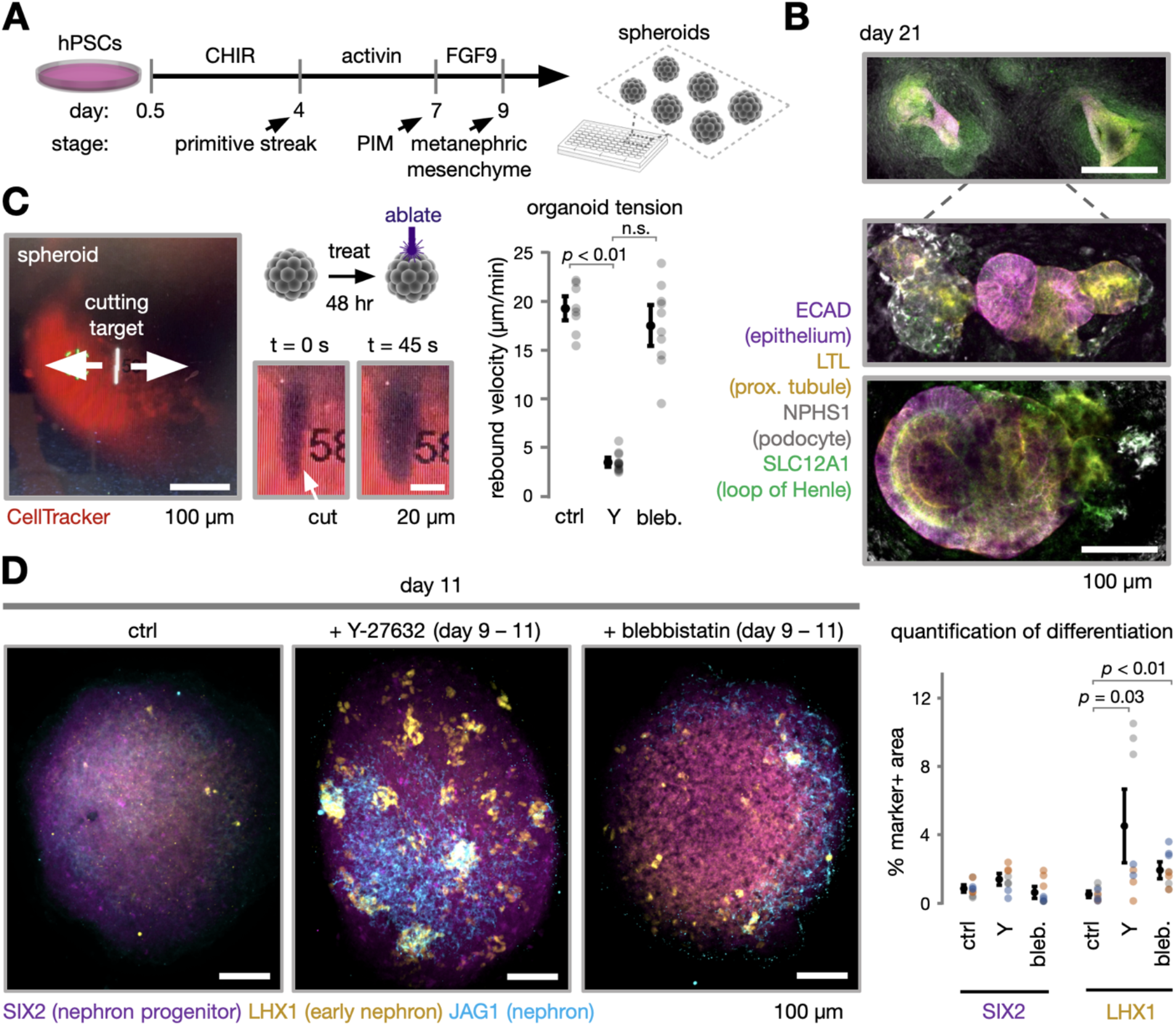
Reducing cell collective tension by ROCK inhibition increases nephron induction in human iPSC-derived kidney organoids. (**A**) Schematic of human iPSC differentiation to metanephric mesenchyme and formation of spheroids for early nephron induction assays. PIM, posterior intermediate mesoderm. (**B**) Confocal immunofluorescence micrograph and insets of iPSC-derived metanephric mesenchyme after continued 2D differentiation in FGF9 basal media until day 21, with a CHIR pulse over days 9-11. (**C**) Laser microdissection of metanephric mesenchyme spheroids. *Left*, Schematic and widefield fluorescence micrographs of example cut location prior to ablation, and rebound of the cut in the 45 s after ablation. *Right*, Plot of cut rebound velocity (change in cut width in 45 s) after ablation for spheroids treated with basal FGF9 media supplemented with CHIR, ± Y-27632 or blebbistatin (mean ± S.D., n > 7 cuts across >3 spheroids per condition. One-way ANOVA, Tukey’s test *p* values indicated). (**D**) *Left*, Representative confocal immunofluorescence projections of organoids after 48 hr perturbations from day 9-11. *Right*, Plot of marker positive pixel areas as a % of projection area for markers of nephron progenitors (SIX2) and early nephron cells (LHX1) (mean ± S.D., *n* > 6 spheroids per condition pooled across three replicates. Replicates are denoted as different colored markers. One-way ANOVA, Tukey’s test *p* values indicated.)

## Discussion

The organ-scale structure of the developing kidney has limited stereotypy between individuals, yet on a smaller scale, its nephron-forming niches exhibit substantial geometric order (Short et al. 2014; Prahl, Viola, et al. 2023). However, the necessity of any particular arrangement of the niche cell compartments for nephron formation is unknown. Our data suggest that the integrity of the cap mesenchyme:stromal boundary is a regulator of nephrogenesis. We perturbed boundary integrity by manipulating cell collective tension via the effects of Rho/ROCK signaling on the actin cytoskeleton, enabling us to explore alternative developmental states of cap mesenchyme:stromal mixing not accessible under homeostatic conditions. We unexpectedly found that increased mixing between these niche compartments in explant cultures correlated with increased nephrogenesis rate, suggesting that developmental regulation of boundary integrity may be a control module for maintaining appropriate renewal:differentiation balance in the niche. This balance is in turn crucial to setting nephron endowment, an important predictor of kidney disease. We went on to narrow interpretation of the mechanism by using primary cell co-culture experiments to find that nephron progenitor-stromal interaction mediates effects of boundary integrity vs. disorder on nephrogenesis.

The mechanism of communication between stromal cells and nephron progenitors dependent on compartment mixing requires further investigation. We expect that ablating especially stromal cell tension with a Rho/ROCK inhibitor reduces the boundary tension:adhesion ratio that is thought to determine boundary integrity (Canty et al. 2017; Heer and Martin 2017; Umetsu et al. 2014). We hypothesize that the resulting increase in heterotypic contact between cell types may lead to increased nephron progenitor exposure to stroma-derived juxtacrine ligands including FAT4, which has been implicated in driving nephron progenitor differentiation (Das et al. 2013; Bagherie-Lachidan et al. 2015). Stromal TGFβ also promotes differentiation of nephron progenitors (Rowan et al. 2018) and multiple barriers may limit the effective diffusion of TGFβ in the niche. Specifically, TGFβ binds to extracellular matrix proteins (Robertson and Rifkin 2016) and stromal cells express TGFβ receptors that may sequester the ligand (Dumbrava et al. 2021). A potentially limited TGFβ diffusion length scale may privilege only those nephron progenitors in close proximity to a stromal cell to receive it.

Our results showed that stromal cells are the most mechanically active niche compartment (defined by traction force and pMLC measurements, **Fig. 1**), suggesting that niche mechanical properties may be dominated by the stromal compartment. Besides effects on biochemical signaling between cell compartments, our data do not rule out a stroma-imposed mechanical microenvironment as a contributor to nephron progenitor decision-making. Indeed, crucial signaling pathways involved in nephrogenesis including Wnt and BMP/SMAD are known to be mechanosensitive in other developmental contexts (Parada et al. 2022; Muncie et al. 2020; Shyer et al. 2017). Similarly, despite some conflicting observations, the effect of FAT4 may be modulated through YAP (Reginensi et al. 2013; Das et al. 2013; McNeill and Reginensi 2017; Bagherie-Lachidan et al. 2015; Mao, Francis-West, and Irvine 2015). Since YAP is a nexus of mechanotransduction signaling (Dupont et al. 2011; Azzolin et al. 2014), there is likely a complex interplay between niche compartment mixing, mechanics, and nephron progenitor differentiation.

What does our perturbation of niche integrity have to do with the native niche under homeostatic conditions? The stroma - cap mesenchyme boundary is dynamic during development (Lawlor et al. 2019). Stromal cells migrate and split the existing cap mesenchyme compartment in two during ureteric bud tip bifurcations (Munro, Hohenstein, and Davies 2017), and nephron progenitors occasionally cross stromal boundaries (Combes et al. 2016). Our recent work has noted variation in cap mesenchyme-stromal compartment mechanical stress (Viola et al. 2023) and mixing over the branching morphogenesis ‘life-cycle’ (Hughes Lab, *in preparation*). Y-27632 is potentially mimicking several of these processes here. The relationship between developmentally programmed variation in niche integrity as a native control mechanism for nephron progenitor renewal versus differentiation is an exciting area for future study.

From an engineering perspective, our data suggest that native niche structure is an important determinant of nephrogenesis rate. This adds emphasis to appropriate control over parameters such as the ratio of nephrogenic niche compartment cell types, their spatial relationship, and their biophysical properties, which are partially tunable through the mechanical microenvironment (Sun, Chen, and Fu 2012). Emerging techniques for high-resolution cell patterning of different cell types in defined microenvironments, combined with a more complete understanding of proximity- and contact-dependent interactions of nephron progenitor niche cell types will inform design of kidney tissue engineering approaches.

## Acknowledgements

We thank Wenli Yang for advice on iPSC maintenance, and Sarah Howden and Melissa Little for advice on iPSC differentiation protocols. We thank Kate Bennett at the Penn Medicine Center for Molecular Studies In Digestive and Liver Diseases, Molecular Pathology and Imaging Core (MPIC, funded under NIH center grant P30-DK050306) for assistance with laser ablation studies. This research was partially supported by the NSF through the University of Pennsylvania Materials Research Science and Engineering Center (MRSEC, DMR-2309043). This work was supported by NIH F32 fellowship DK126385 (LSP), the Predoctoral Training Program in Developmental Biology T32HD083185 (JMV), NSF GRFP (JL), NIH NIGMS MIRA R35GM133380 (AJH), NIH NIDDK R01DK132296 (AJH), and a Penn Center for Precision Engineering for Health (CPE4H) pilot grant (AJH).

## Supplementary Information

### Supplementary Figures

**Fig S1:**
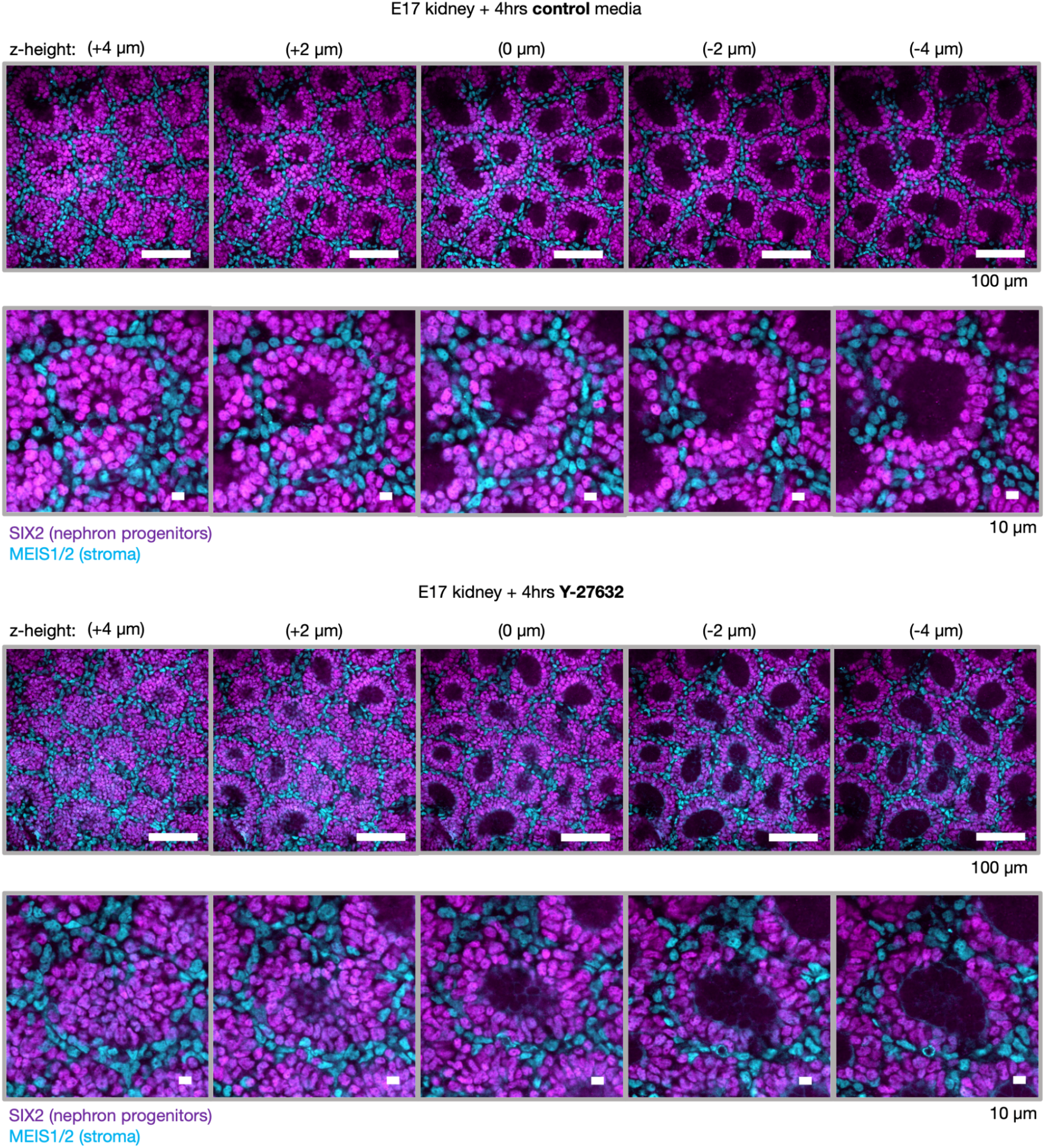
Y-27632 treatment increases nephron progenitor-stromal mixing across optical planes spanning the cap mesenchyme. *Top*, confocal whole-mount immunofluorescence micrographs and insets of E17 mouse embryonic kidney explants after 4 hr culture for a representative field of view. Columns are images at different *z*-heights through the niche. *Bottom*, similar images after 4 hr ROCK inhibitor treatment.

**Fig. S2:**
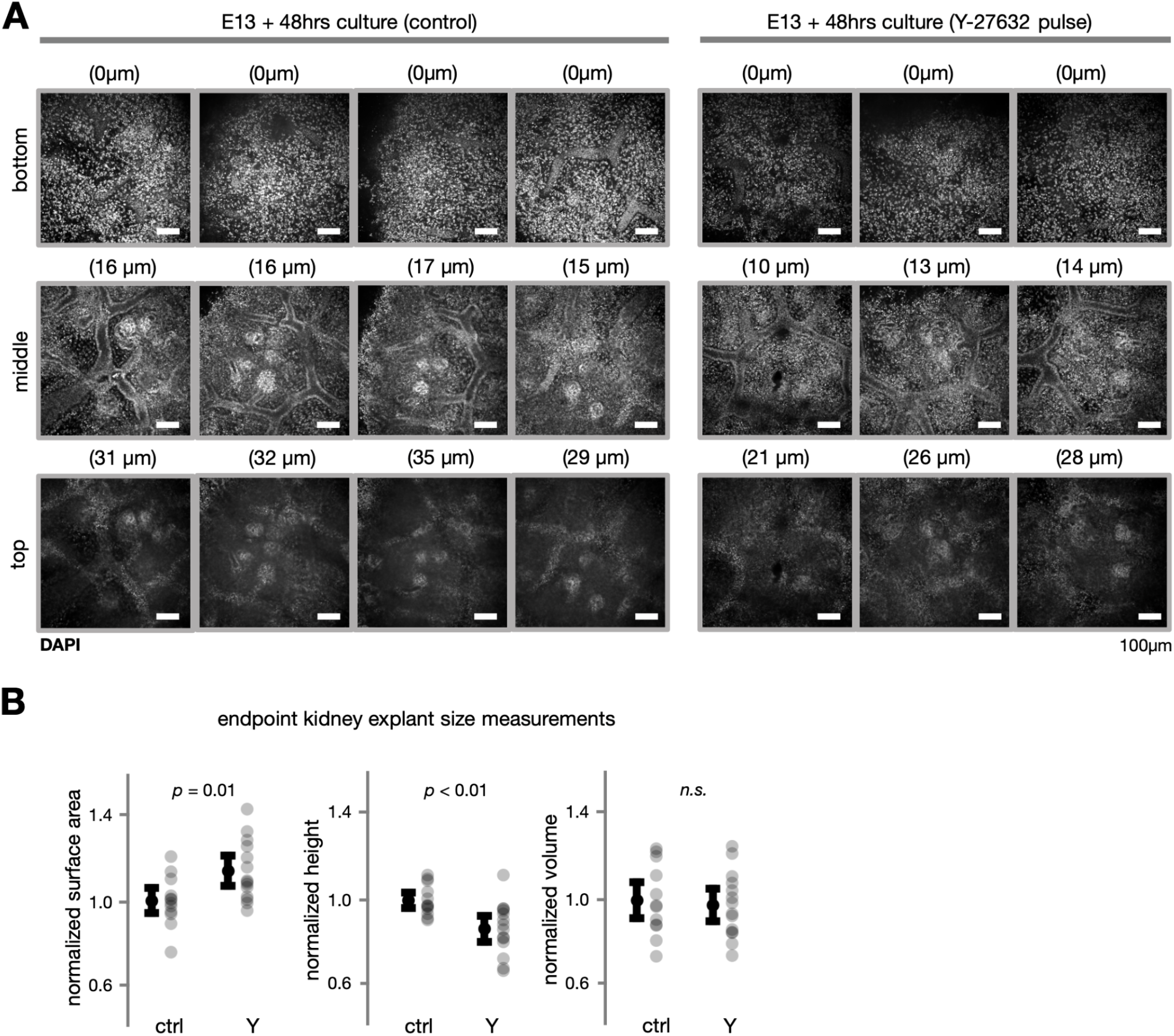
Transient Y-27632 treatment does not affect average kidney volume in explant culture. (**A**) Semi-quantitative scoring of kidney explant bottom, middle, and top planes by confocal immunofluorescence microscopy after 48 hr culture ± Y-27632. (**B**) Plots of size metrics after culture period.

**Fig. S3:**
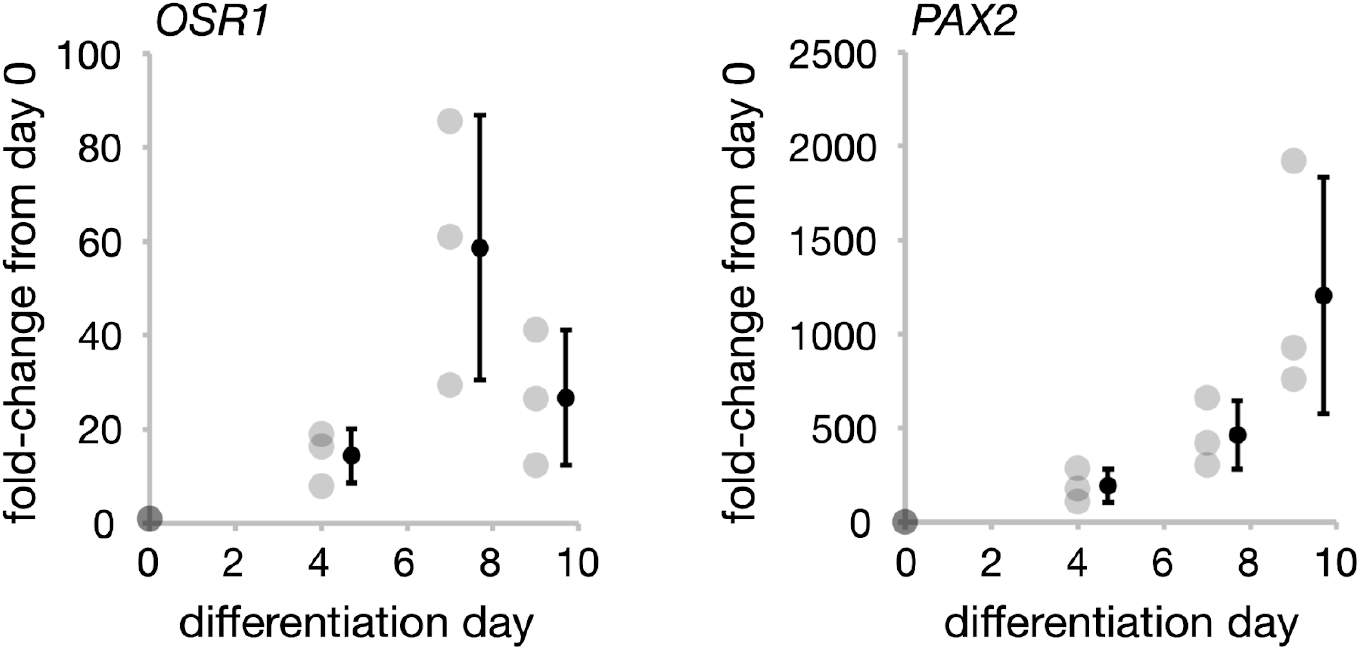
qPCR validates human iPSC differentiation through posterior intermediate mesoderm to metanephric mesenchyme. Plots of transcript fold-change in the respective markers for these stages *OSR1* and *PAX2* (see Methods) for human iPSCs differentiated over days 0-9 according to the Morizane protocol (Morizane and Bonventre 2017) (mean ± S.D., *n* = 3 wells per condition).

## Methods

### Animal experiments

Mouse protocols followed NIH guidelines and were approved by the Institutional Animal Care and Use Committee of the University of Pennsylvania. E14-E17 embryos were collected from timed pregnant CD-1 mice (Charles River) and stages confirmed by limb anatomy as previously described (Wanek et al. 1989). Embryonic kidneys were dissected in chilled Dulbecco’s phosphate-buffered saline (DPBS, MT21-31-CV, Corning) (Barak and Boyle 2011).

### E17 embryonic kidney short term Y-27632 incubation

E17 mouse embryonic kidneys were dissected and kept in DPBS in 1.7 ml tubes. Kidneys were placed in a 1% agarose-coated well of a 24-well tissue-culture plate to prevent them from adhering to the substrate. 250 µL of Dulbecco’s Modified Eagle Medium (DMEM) (Corning, 10-013-CM) + 5% fetal bovine serum + 1% Pen/Strep (Mediatech, MT30-002-CI) +/- 30 µM Y27632 (StemCell Technologies, 72304) was added to just cover the dissected kidneys. Kidneys were then incubated for 4 hr in a standard 5% CO_2_ tissue culture incubator at 37°C and subsequently fixed in ice-cold 4% paraformaldehyde in DPBS for 20 min (see **Kidney immunofluorescence imaging**).

### Cap mesenchyme and stromal compartment geometric analysis

The sum of two Gaussian distributions was fitted to SIX2 intensity profiles across each stromal gap in MATLAB. The full width at half maximum was calculated for both Gaussian distributions, representing the width of each cap mesenchyme compartment on either side of the gap. Width measurements derived from qualitatively poor Gaussian fits were excluded from analysis. The PDGFRA immunofluorescence channel was used to manually annotate the stromal region. 200 pixel background subtraction was used for the SIX2 channel in FIJI. SIX2+ nuclei were manually thresholded in each individual image. Pixel overlap was normalized within litters before plotting.

### Embryonic kidney explant culture

Explant culture was adapted from previous methods (Costantini et al. 2011). Dissected E13 mouse embryonic kidneys were transferred to ice-cold PBS in 1.7 ml microcentrifuge tubes. Kidneys were then placed onto transwell filter inserts (Corning prod. # 3413) in 12-well tissue culture plates. Pipette tips were cut to a diameter larger than kidneys with a razor blade and coated with 1% gelatin in DPBS. Kidneys were transferred with these tips to avoid organ adhesion to the tip walls. Excess DPBS surrounding the kidney was removed from the filter. 250 µL of media per well was pipetted into underneath transwell filters. Kidneys were cultured in a standard 5% CO_2_ tissue culture incubator at 37°C and media was changed at the indicated time points. At culture endpoint kidneys were fixed in ice-cold 4% paraformaldehyde in DPBS for 20 min. After fixation, transwell filters were cut out with a scalpel and filter-adhered kidneys were further processed for immunofluorescence microscopy (see **Kidney immunofluorescence imaging**). For imaging, transwell filters with adhered kidneys were placed against the bottom of a cover slip-bottom tissue culture plate (Greiner 662892).

### Quantitation of endpoint explant height and volume

Kidney volume was calculated by multiplying endpoint kidney surface area and height. Surface area was calculated by manual segmentation of endpoint brightfield images of kidney explants. Kidney height was quantified by estimating top and bottom confocal immunofluorescence slices from the DAPI channel at 20x using 1 µm *z*-step size at the center of each explant (**Fig. S2**).

### Embryonic kidney explant culture endpoint JAG1+ nephron structure annotation

Explant kidneys were imaged using 20X confocal *z* stacks with 5 µm step size spanning whole kidney explants. Distinct JAG1+ structures were manually annotated by examining individual image stacks. 10x images of whole kidney explants were cross-referenced so that nephron structures that spanned across multiple 20x stacks were counted only once.

### Primary mouse embryonic kidney nephrogenic whole-niche and nephron progenitor-enriched spheroids

Primary embryonic mouse nephrogenic whole niche spheroids were made by adapting previously described protocols (Brown et al. 2013, 2011; Brown, Muthukrishnan, and Oxburgh 2015). Briefly, a digest solution consisting of 0.5% pancreatin (Sigma-Aldrich, P7545) and 0.25% Collagenase A (Sigma-Aldrich, C0130) in DPBS was made. Pancreatin was first added to DPBS and allowed to solubilize on a rotator at 4°C overnight. Collagenase was added to the digest solution and the mixture was solubilized on a rotator at room temperature for 3-5 hr on the day of the experiment. Batches of 8 dissected E17 embryonic kidneys were incubated in 1.5 ml digest solution for 15 min at 37°C. The digest reaction was then stopped by adding 75 µL of fetal bovine serum. 1.4 ml of the digest solution containing dissociated cells was then transferred to a fresh microcentrifuge tube containing 2 µL of DNase (Fisher Scientific, AM2238) and incubated for 10 min at 37°C. The cell solution was centrifuged and resuspended 2 times in 1 ml Hank’s Balanced Salt Solution (HBSS) + 5% fetal bovine serum. Cells were then resuspended in media and passed twice through 40 µm cell strainers by pipetting. Cells were plated at 50,000 cells per well in 96-well ultra low attachment round-bottom plates (Corning 7007) in Advanced RPMI 1640 Medium, Glutamax, 1x Penn/Strep and 10 ng ml^-1^ FGF9 (‘basal media’) plus the specified perturbation conditions, which included 3 µM CHIR 99021 (Tocris, 4423) +/- 10 µM Y27632 (StemCell Technologies, 72304) from a 10 mM stock in DMSO. Cells were then aggregated by plate centrifugation at 300 g for 30 s and transferred to a standard tissue culture incubator. At experiment endpoints, spheroids were collected and transferred to 1.7 ml tubes using a p200 pipette tip widened by cutting the tip to size with a razor blade. Excess media was removed and spheroids were fixed in ice-cold 4% paraformaldehyde in PBS for 15 min. Spheroids were then stained for immunofluorescence microscopy similar to kidneys (see **Kidney immunofluorescence imaging.**)

Nephron progenitor-enriched spheroids were made by removing unwanted cell compartments by negative sorting of whole niche cell mixtures using benchtop magnetic cell sorting. After whole-niche cell mixtures were collected, a solution of biotinylated primary antibodies in DPBS + 3% fetal bovine serum + 10mM EDTA (Invitrogen, 15575-038) was prepared. Primary antibodies and dilutions are as follows: anti-CD326 (1:100, 13-5791-80, ThermoFisher, RRID: AB_1659715), anti-TER-119 (1:50, ThermoFisher, 13-5921-81, RRID:AB_466796), anti-CD105 (1:50, ThermoFisher, 13-1051-81, RRID:AB_466555), and anti-CD140A (1:25, ThermoFisher, 13-1401-80, RRID:AB_466606). Cells were suspended in DPBS + 3% fetal bovine serum + 10mM EDTA at 2 x 10^7^ cells/ml. An equal amount of previously prepared 2X primary antibody solution was added to the cell mixture and mixed well by pipetting. The mixture was incubated for 10 min at room temperature. The volume was then brought up to 4 ml with DPBS + 3% fetal bovine serum + 10 mM EDTA and mixed by pipetting. The mixture was centrifuged at 300 g for 5 min and cells were resuspended at 10^7^ cells/ml. Streptavidin-linked magnetic beads (ThermoFisher, MSNB-6002-71) were added to the cell mixture at 1:10, mixed well by pipetting, and the cell solution was incubated at room temperature for 5 min. The cell solution was brought up to 2.5 ml with DPBS + 3% fetal bovine serum + 10 mM EDTA and aliquoted into two 1.5 ml microcentrifuge tubes. Tubes were placed on a magnetic tube holder (Sergi Lab Supplies, B0812XLPVK) and incubated for 5 min at room temperature. The tube rack was inverted and the cell mixture poured into a collection tube so that negatively sorted cells were left behind. The collected cell mixture was placed in fresh tubes and placed on the magnetic rack and the latter two steps were repeated. The collected enriched nephron progenitor cell mixture was resuspended in Advanced RPMI 1640 Medium, Glutamax, 1x Penn/Strep and 10 ng ml^-1^ FGF9 (‘basal media’) and spheroid generation and further experimental procedures were carried out as in whole-niche spheroids.

### Kidney immunofluorescence imaging

mmunofluorescence staining and imaging was performed as previously described (Prahl et al. 2021), using protocols adapted from Combes *et al*. and O’Brien *et al*. (Combes et al. 2014; O’Brien et al. 2018). Briefly, dissected kidneys were fixed in ice cold 4% paraformaldehyde in DPBS for 20 min, washed three times for 5 min per wash in ice cold DPBS, blocked for 2 hr at room temperature in PBSTX (DPBS + 0.1% Triton X-100) containing 5% donkey serum (D9663, Sigma), incubated in primary and then secondary antibodies in blocking buffer for at least 48 hr at 4°C, alternating with 3 washes in PBSTX totaling 12-24 hr. The minimum duration of primary and secondary incubations and washes depended on the age of the kidney, as previously described (Combes et al. 2014). Primary antibodies and dilutions included rabbit anti-Six2 (1:600, 11562-1-AP, Proteintech, RRID: AB_2189084), goat anti-E-cadherin (1:200, AF748, R&D systems, RRID: AB_355568), rat anti-PDGFRA (1:200, 14-1401-82, Invitrogen, AB_46749), goat anti-ITGA8 (1:20, AF4076, R&D systems, RRID: AB_2296280), rabbit anti-LEF1 (1:200, 2230, Cell Signaling Technologies, RRID: AB_823558), rat anti-E-cadherin (1:100, ab11512, abcam, RRID:AB_298118), mouse anti-E-cadherin (1:200, clone 34, 610404, BD Biosciences, RRID: AB_397787), rabbit anti-MYL12A (phospho S19) (1:100, ab2480, abcam, RRID:AB_303094), goat anti-jagged 1 (1:150, AF599, R&D Systems, RRID: AB_2128257), rabbit anti-RET (1:200, 3223, Cell Signaling Technologies, RRID:AB_2238465) Secondary antibodies (all raised in donkey) were used at 1:300 dilution and include anti-rabbit AlexaFluor 405 (ThermoFisher, A48258, RRID: AB_2890547), anti-rat AlexaFluor 555 (ThermoFisher, A48270, RRID: AB_2896336), anti-goat AlexaFluor 488 (A11055, ThermoFisher, RRID: AB_2534102), anti-rat AlexaFluor Plus 405 (A48268, ThermoFisher, RRID: AB_2890549), anti-rabbit AlexaFluor 555 (A31570, ThermoFisher, RRID: AB_2536180), anti-goat AlexaFluor 647 (A32849, ThermoFisher, AB_2762840), anti-mouse AlexaFluor 405 (A48257, ThermoFisher, RRID: AB_2884884), anti-rabbit AlexaFluor 488 (A21206, ThermoFisher, RRID: AB_2535792). In some experiments, samples were counterstained in 300 nM DAPI (4’,6-diamidino-2-phenylindole; D1306, ThermoFisher) diluted in blocking buffer for 2 hr at room temperature, followed by 3 washes in PBS.

Kidneys were imaged in wells created with a 2 mm diameter biopsy punch in a ∼5 mm-thick layer of 15:1 (base:crosslinker) polydimethylsiloxane (PDMS) elastomer (Sylgard 184, 2065622, Ellsworth Adhesives) set in 35 mm coverslip-bottom dishes (FD35-100, World Precision Instruments). Imaging was performed using a Nikon Ti2-E microscope equipped with a CSU-W1 spinning disk (Yokogawa), a white light LED, laser illumination (100 mW 405, 488, and 561 nm lasers and a 75 mW 640 nm laser), a Prime 95B back-illuminated sCMOS camera (Photometrics), motorized stage, 4x/0.2 NA, 10x/0.25 NA and 20x/0.5 NA lenses (Nikon), and a stagetop environmental enclosure (OkoLabs).

### Human iPSC-derived nephron progenitors

Nephron progenitor cells were generated from SIX2^EGFP^ transgenic reporter iPSC line (*SIX2*-T2A-*EGFP*, Murdoch Children’s Research Institute / Kidney Translational Research Center, Washington University Nephrology) (Vanslambrouck et al. 2019) according to published protocols (Morizane and Bonventre 2017). Briefly, iPSCs were maintained in standard tissue-culture treated 24-well plates in stem cell maintenance medium plus supplement (mTeSR+ kit, StemCell Technologies 100-0276) and 1x Penn/Strep from a 100x stock (Mediatech, MT30-002-Cl). Cells were passaged using Accutase (StemCell Technologies, 07920) and plated at a density of 25,000 cells per well. Differentiation was induced with Advanced RPMI 1640 Medium (Fisher Scientific, 12-633-012), GlutaMAX (ThermoFisher, 35050061), 1x Penn/Strep and 7µM CHIR 99021 (Tocris, 4423) either the next day or when cells reached 50-75% confluency. On day 4, media was changed to Advanced RPMI 1640 Medium, Glutamax, 1x Penn/Strep and 10 ng ml^-1^ activin (R&D Systems, 338AC010CF). On Day 7 media was changed to Advanced RPMI 1640 Medium, Glutamax, 1x Penn/Strep and 10 ng ml^-1^ FGF9 (R&D Systems, 273-F9-025). On day 9 (metanephric mesenchyme stage), cells were either 1) differentiated further in 10 ng ml^-1^ FGF9 until day 21 with a 3 µM CHIR pulse over days 9-11, 2) dissociated and aggregated for use in spheroid experiments, or 3) cultured for up to 3 days in 10 ng ml^-1^ FGF9 media to maintain them in the SIX2+ state.

### Validation of iPSC differentiation by qPCR

RNA extraction was performed on cells from individual wells of a 24-well tissue-culture dish at each differentiation time-point using an RNEasy mini kit (Qiagen 74104). cDNA was generated from RNA using an Applied Biosystems High Capacity cDNA Reverse Transcription kit (ThermoFisher 4368814), following the manufacturer protocol and using 2 ng of RNA in the reaction for each sample. Applied Biosystems Sybr Green PCR Master Mix (ThermoFisher 4344463) was used for qPCR in an Applied Biosystems 7300 thermocycler. The Applied Biosystems protocol for Sybr Green PCR Master Mix was followed for thermocycler temperatures and times. 25 ng of cDNA was used in each reaction along with the appropriate qPCR primers at 300 nM. qPCR primers for *ACTB*, *OSR1*, and *PAX2* were used as previously described (Morizane et al. 2015). DeltaDeltaCt values were calculated using *ACTB* as a housekeeping gene.

### iPSC-derived nephron organoid differentiation assays

Day 9 metanephric mesenchyme-stage nephron progenitors were lifted with Accutase and re-plated at 50,000 cells per well in 96-well ultra low attachment round-bottom plates (Corning 7007) in Advanced RPMI 1640 Medium, Glutamax, 1x Penn/Strep and 10 ng ml^-1^ FGF9 (‘basal media’) plus the specified perturbation conditions, which included 3 µM CHIR, 10 µM Y27632 (StemCell Technologies, 72304) from a 10 mM stock in DMSO, and/or 30 µM (S)-(-)-blebbistatin (Tocris, 1852) from a 30 mM stock in DMSO. Cells were then aggregated by plate centrifugation at 300 g for 30 s. After 48 hr in culture, spheroids were collected and transferred to 1.7 ml tubes using a p200 pipette tip widened by cutting the tip to size with a razor blade. Excess media was removed and spheroids were fixed in ice-cold 4% paraformaldehyde in PBS for 15 min. Spheroids were then stained for immunofluorescence microscopy similar to kidneys (see **Kidney immunofluorescence imaging.**) via primary and secondary antibody incubation for 48 hr each at 4°C and washes in PBSTX totaling 6 hr. Primary antibodies and dilutions included mouse anti-LHX1 (1:50, 4F2, Developmental Studies Hybridoma Bank, University of Iowa, RRID: AB_531784) and rabbit anti-SIX2 (1:600, 11562-1-AP, Proteintech, RRID: AB_2189084). Spheroids were then incubated in 1 µg ml^-1^ DAPI in PBS at 4°C for 1hr before washing once in ice-cold PBS. Spheroids were transferred for imaging using a widened p200 pipette tip to a cover-slip bottom 24-well plate (Greiner 662892). Excess PBS was removed and spheroids were incubated with ∼20 µl of FocusClear (Cedarlane Labs, FC-101) for 20 min at room temperature prior to confocal imaging. Spheroids were imaged using 5 µm z-steps.

### Quantification of primary mouse cell and iPSC-derived spheroid cell differentiation

Maximum intensity projections of spheroids were generated and further processed in FIJI. DAPI signal or manual annotation was used to segment spheroids and calculate their areas. SIX2 and LHX1 signal was segmented using 200 pixel rolling-ball background subtraction and thresholded to remove background signal. The total numbers of SIX2+ and LHX1+ pixels were then divided by the total number of DAPI+ pixels to normalize positive pixels by spheroid area.

### Laser ablation

Nephron organoids were labeled in 1:1000 CellTracker Red (ThermoFisher C34552, 10 mM stock in DMSO) in Dulbecco’s minimum essential medium (DMEM, 10-013-CV, Corning) for 1 hr at 37°C, washed 3x in DMEM and placed in 2mm-diameter PDMS wells (see **Kidney immunofluorescence imaging**), this time plasma bonded to a quartz coverslip (No. 1 thickness, Ted Pella, 26014) to maximize UV transmission. Organoids were ablated using a 355 nm UV laser (Molecular Machines & Industries, SL µCut v1.0) through a 10x objective on a Nikon Ti2 controlled by MMI software. Cut opening time-lapses were imaged from the microscope monitor using an iPhone 8 video camera and spatially calibrated using on-screen fiducial marks in the MMI software.

### Traction force microscopy

A 100 mm-diameter mechanical-grade silicon wafer (University Wafer) was fabricated with shims for precise gel spacing as previously described (Viola et al. 2020). Briefly, a ‘shim’ photomask reflecting the geometry of a standard 1’’ x 3’’ microscope slide (12-559-A3, ThermoFisher) was printed at 20,000 d.p.i (CAD/Art Services) to cast parallel rails on the wafer to allow 30 µm-thick gel manufacturing later. The wafer was first baked on a hot plate at 200°C for 10 min and then centered on a vacuum spin coater (SCK-300P, Instras Scientific). Microchem SU-8 2025 photoresist (Fisher Scientific) was dispensed on the wafer at 4 ml and spin-coated at 500 rpm for 12 s followed by 3,000 rpm for 45 s to create a uniform 30 µm layer of photoresist. The assembly was then baked on a hot plate at 95°C for 8 min. The photomask was placed on the treated wafer and exposed to 365 nm ultraviolet light at 175 mJ/cm^2^ (M365LP1, Thorlabs). It was further baked at 95°C for 8 min and developed by submerging in SU-8 developer (Microchem SU-8 developer, Fisher Scientific) for at least 15 min. The wafer was rinsed with acetone and isopropanol and left to air dry. Vapor deposition was used to render the wafer hydrophobic with 1 ml of dichlorodimenthysilane (DCDMS, Sigma).

Standard microscope slides were functionalized for polyacrylamide (PA) gel covalent attachment by etching in 1N sodium hydroxide (NaOH, S8045, Sigma) for 15 min and silanizing with 3-(trimethoxysilyl)propyl methacrylate (440159, Sigma), acetic acid (EMD Millipore), and deionized (DI) water at 2:3:5 ratio by sandwiching 150 µL solution between two slides. The ‘slide sandwiches’ were incubated for at least 30 min at room temperature and rinsed with 100% methanol and DI water.

The silanized side of the treated slide was placed face down on top of the shim on the wafer. PA gel precursor solution was prepared by pipetting 645.5 µL DI water, 187.5 µL acrylamide solution (40% w/v Acrylamide Solution, 1610140, Bio-Rad), 100 µl 10x DPBS (14200075, Thermo Fisher Scientific), 35 µL N,N-methylenebisacrylamide crosslinker (2% w/v bis-acrylamide Solution, 1610142, Bio-Rad), 20 µL of 1:100 diluted fluorescent beads in DI water (FluoSpheres carboxylate-modified 0.2 µm, 660/680, F8807, Thermo Fisher Scientific), 6 µL 10% v/v tetramethylethylenediamine (TEMED) (Sigma) and 6 µL 10% w/v ammonium persulfate (Sigma) in DI water. PA gels formed from this precursor have a Young’s modulus of 6 kPa (Lakins, Chin, and Weaver 2012). The precursor was degassed under vacuum in an ultrasonic cleaning bath (15-337-411, Thermo Fisher Scientific), and 120 µL was dispensed in the gap between the slide and wafer and allowed to cure for 30 min protected from light exposure.

To derivatize uncoupled acrylamide chains on the gel surface with NHS ester for adhesive protein functionalization, 0.01% (w/v) N,N-methylenebisacrylamide in 50 mM (pH 6.0) 4-(2-hydroxyethyl)-1-piperazineethanesulfonic acid (HEPES, 40820004, bioWORLD) buffer was mixed with 0.09% (w/v) lithium phenyl-2,4,6-trimethylbenzoylphosphinate (LAP photoinitiator, 900889, Sigma) and 0.1% (w/v) acrylic acid-*N*-hydroxysuccinimide ester (N2 acrylic acid, A8060, Sigma), as previously reported (Prahl, Porter, et al. 2023). Free-radical photochemistry was carried out by pipetting 200 µl of the mixture on top of the PA gel and initiated under UV-light (I365 nm = 15 mW cm^−2^) for 30 min. The NHS-coated PA gel was washed in ice-cold DI water and 100 µM sodium chloride (NaCl, S6191, Sigma) and assembled into a clip-on 8-well chambered slide with a silicone gasket (CCS-8, MatTek). The wells were incubated in 100 mg/ml Matrigel (356234, Corning) overnight at 4°C to covalently couple the N-terminus of the ECM proteins before washing slides four times with PBS at room temperature. After the last wash, primary cell culture media was used to incubate the wells before cell seeding.

E17 kidneys were dissected, dissociated (see **Primary mouse embryonic kidney nephrogenic whole-niche and nephron progenitor-enriched spheroids**), and passed through a 40 µm cell strainer (CLS431750, Corning). Cells were seeded at 20,000 cells per well and cultured for 12 hr to allow attachment and traction equilibration. Brightfield and Cy5 confocal images of cell and bead positions were then captured in the full-traction state. An automated *xy* point assignment program was populated to record user-defined multipoints within each well, and autofocus was used to automate collection of *z*-stacks.

To read out the traction forces, cells were fixed in 4% paraformaldehyde (16% PFA, 15710, Electron Microscopy Sciences) in 1x DPBS for 20 min and washed with DPBS four times at 5 min each. The wells were loaded on the confocal microscope and global registration performed by manually fine-tuning the four corners of the slide to align the same positions previously recorded in the full-traction state. After global translational and rotational displacements were corrected, the same *xy* points were imaged. Autofocus was used to capture *z*-stacks of the beads identical to those of the full-traction state.

For downstream immunofluorescence, cells were permeabilized in 0.5% v/v Triton X-100 for 5 min in 1x DPBS and blocked with 1% BSA in IF wash buffer (1% w/v BSA, 2% v/v Triton X-100; 0.41% v/v Tween-20, EMD Millipore, 1x DPBS) for 1 hr. The wells were then incubated with fluorophore-conjugated primary antibodies: goat anti-RET-647 (1:50, AF1485, Bio-Techne, RRID: AB_354820), rat anti-PDGFRA-555 (1:200, 14-1401-82, Invitrogen, RRID: AB_46749), and rabbit anti-SIX2-488 (1:200, 11562-1-AP, Proteintech, RRID: AB_2189084). Primary antibodies were conjugated by clean up with AbSelect (861-0010, Novus) and labeling with Lightning-Link Antibody Labeling Kits (336-0010, 333-0010, 332-0010, Novus) following user manuals. Cells were washed with IF buffer 3 x 1 hr each while protected from light, and nuclei were counterstained with 150 nM DAPI for 15 min. Finally, wells were washed with IF buffer and PBS for 1 hr each before confocal immunofluorescence imaging of antibody markers using 20 µm *z*-stacks above the gel surface after a similar *xy* registration procedure.

Image analysis was performed using automated macros in ImageJ/Fiji (Schindelin et al. 2012). “Find Focused Slides” were used to select the corresponding *z*-frames between the full-traction and relaxed states, and z-stack maximum projections were concatenated to run registration with the “Linear Stack Alignment with SIFT” plugin (Lowe 2004). Displacement and traction fields were calculated with the PIV and FTTC plugins respectively (Tseng et al. 2012). Automated cell segmentation was carried out with the StarDist plugin (Schmidt et al. 2018) and only single cells were analyzed. Specifically, the DAPI channel was used to segment cell nuclei, and cells were excluded if their diameters were smaller than 5 µm or larger than 20 µm. If the Euclidean distance between two nuclei was smaller than twice the sum of their major axes, both cells were excluded. The filtered ROIs were then looped through to extract the mechanical and biomolecular information of the cells. A 60 x 60 µm window centered on each cell was drawn to calculate the average traction force and pixel intensity, counting the top 20th and 10th percentiles of the pixels, respectively. The cell ID number along with its traction and IF data was stored before further processing. For cell identification, 50 positive cells of a specific cell type were manually selected based on the known marker expression pattern, and the signals from the channel corresponding to the positive marker were pooled. Fluorescence cutoffs were determined by taking the 5th percentile of this population. Thresholds of positive cell types were used to assign cell identities. If a cell had a marker expression level higher than the positive cutoff, it was assigned to the corresponding cell type. If a cell had expression levels higher than all cutoffs, it was taken to be cell debris and excluded from the analysis. Less than 0.1% of such artifacts were observed in this study. Finally, the average traction force of different cell types were calculated.

### Plotting and statistical analysis

One-way analysis of variance (ANOVA) with correction for multiple comparisons using Tukey’s honestly significant difference test was performed in MATLAB using anova1.m and multcompare.m functions. Paired and unpaired *t*-tests were also performed in MATLAB. Violin plots were generated in MATLAB using the violinplot-MATLAB plugin (Bastian Bechtold).

